# A Neural Index Reflecting the Amount of Cognitive Resources Available during Memory Encoding: A Model-based Approach

**DOI:** 10.1101/2022.08.16.504058

**Authors:** Si Ma, Vencislav Popov, Qiong Zhang

## Abstract

Humans have a limited amount of cognitive resources to process various cognitive operations at a given moment. The Source of Activation Confusion (SAC) model of episodic memory proposes that resources are consumed during each processing and once depleted they need time to recover gradually. This has been supported by a series of behavioral findings in the past. However, the neural substrate of the resources is not known. In the present study, over an existing EEG dataset of a free recall task (Kahana et al., 2022), we provided a neural index reflecting the amount of cognitive resources available for forming new memory traces. Unique to our approach, we obtained the neural index not through correlating neural patterns with behavior outcomes or experimental conditions, but by demonstrating its alignment with a latent quantity of cognitive resources inferred from the SAC model. In addition, we showed that the identified neural index can be used to propose novel hypothesis regarding other long-term memory phenomena. Specifically, we found that according to the neural index, neural encoding patterns for subsequently recalled items correspond to greater available cognitive resources compared with that for subsequently unrecalled items. This provides a mechanistic account for the long-established subsequent memory effects (SMEs, i.e. differential neural encoding patterns between subsequently recalled versus subsequently unrecalled items), which has been previously associated with attention, fatigue and properties of the stimuli.

## A Neural Index Reflecting the Amount of Cognitive Resources Available during Memory Encoding: A Model-based Approach

Humans are only able to process and maintain a relatively small amount of information at any given moment and one explanation, common to many models of working memory, is that processing depends on a limited amount of cognitive resources (Anderson et al., 1996; Ma et al., 2014; Oberauer et al., 2016). Even though, in contrast, long-term memory is often considered to have unlimited capacity, at least some theories assume that the rate of encoding in long-term memory depends on the amount of working memory resources available at the time of encoding (Atkinson & Shiffrin, 1968; Popov & Reder, 2020; Reder et al., 2007). For example, the Source of Activation Confusion (SAC) model of episodic memory (Popov & Reder, 2020; Reder et al., 2000; Reder et al., 2007) assumes that each time an item is stored in long-term memory (LTM), its encoding depletes a fixed proportion of the currently available resources and that the strength of the resulting memory trace is proportional to the amount of resources assigned to it. The amount of resources is finite, and once depleted it needs time to recover gradually (i.e. resource-depletion-and-recovery assumption). This resource assumption has been supported recently by numerous behavioral findings (Kowialiewski et al., 2021; Mizrak & Oberauer, 2021; Oberauer, 2022; Popov et al., 2019; Popov & Reder, 2020; Popov et al., 2021). Despite the behavioral evidence, this resource-depletion-and-recovery assumption has never been tested on neural data and we currently do not know what the biological or neural substrate is of the proposed limited resources.

In this paper, we establish a neural signature indexing the amount of resource available, providing further support for the proposed limited resources. To **determine such a neural index of resource availability**, we used a free recall paradigm where participants were asked to remember a list of consecutively presented words and later to recall all the words from the list in any order. We used the power spectrum pattern of the EEG signal during the encoding of a word as the key neural signature to derive the amount of available cognitive resources. Consistent with SAC’s resource-depletion-and-recovery assumption, we assume that the available cognitive resources are at their fullest at the beginning of each list. We therefore defined the power spectrum pattern of encoding the first word in the list as a template pattern for maximum resources. We then measured how much the encoding patterns of other words are similar to this template pattern. We used the *pattern similarity* as an index of resource availability - we reasoned that the more a pattern is similar to that of the maximum possible resources, i.e. neural pattern at the beginning of each list, the more resources are available and the higher the resource availability index value will be.

To **validate the neural index of resource availability**, we used a model-based approach. We fit the SAC model to two recall behavioral findings that according to the model can be explained by differential resource consumption during encoding - the primacy serial position effect (Murdock, 1962; Oberauer, 2022) and the sequential effect of word frequency (Popov & Reder, 2020). Based on the fit to the recall data, the model infers the amount of resources that are available during the encoding of each word on the list. If the identified neural signature represents resource availability, it should index an amount of available cognitive resources consistent with that inferred from the recall performance using the SAC model. This will both validate the neural index of resource availability and provide a strong test of the SAC’s resource-depletion-and-recovery assumption as a different portion of the data is used for deriving the neural index (i.e. neural data during the encoding phase) than for fitting the SAC model (i.e. behavioral data during the recall phase). The two relevant behavioral effects are:

1. *Primacy serial position effect* - a key aspect of delayed free recall is the primacy effect - recall probability is often highest for the first presented item and it decreases gradually with the serial position until it asymptotes in the middle of the list (Murdock, 1962). Previous fits of the SAC model have shown that the resource-depletion-and-recovery mechanism can account not only for the main effect of primacy (Popov & Reder, 2020), but also the fact that the primacy effect is steeper with faster presentation rates (Oberauer, 2022) and with low-frequency words (Popov & Reder, 2020). According to the SAC model, the primacy effect occurs because the amount of available resources decreases as cognitive operations move forward. In a free recall task, participants have their full pool of cognitive resources at the beginning of a list. Encoding the first word depletes a portion of the resource pool. Since resources recover only gradually over time, participants will have a partially depleted resource pool when they start to encode the second word and such depletion accumulates as the list moves forward. Eventually, the proportion of resources depleted by each word and the amount recovered between words balance each other, leading to an asymptote of available resources, mirroring the behavioral asymptote. We anticipate such accumulation of depletion will be reflected by the changes in the neural index of resource availability, i.e. the neural index decreases as the list continuously moves forward until it reaches the asymptote.
2. *Sequential effect of word frequency* - According to the SAC model, based on prior work (Diana & Reder, 2005; Reder et al., 2007), the encoding of low-frequency words requires more resources than the encoding of high-frequency words, leaving fewer resources to process subsequent information and therefore impairing memory performance (for a review, see Popov and Reder, In press). Because the encoding of low-frequency words depletes more resources relative to high-frequency words, the model predicts a sequential effect of word frequency in episodic memory - namely that memory performance for one word in a study list should be higher when it follows high-frequency rather than low-frequency words. In support of this prediction, Popov and Reder (2020) discovered that the word frequency of a specific word, *X*_*k*_, where k represents the word’s position on a study list, affected memory for the items that followed it on the study list, i.e. items *X*_*k*+1_, *X*_*k*+2_, etc. Specifically, memory performance was better across a variety of test types (item and associative recognition, cued and free recall) for item *X*_*k*+1_ when the preceding *X*_*k*_ item was a high-frequency word rather than a low-frequency word. If the identified neural index in fact represents resource availability, it should measure a lower amount of cognitive resources during encoding *X*_*k*+1_ after encoding a low-frequency word *X*_*k*_ compared to a high-frequency word *X*_*k*_. Moreover, the SAC model can also capture a stronger effect of word frequency on the memory of the immediate subsequent item *X*_*k*+1_ compared with the next subsequent item *X*_*k*+2_ since the resources have less time to recover by the time an item is presented for *X*_*k*+1_ than *X*_*k*+2_ (Popov & Reder, 2020). Therefore, a valid neural index of resource availability should index a larger difference at *X*_*k*+1_ compared with *X*_*k*+2_ between the resource availability following the encoding of a low-frequency word and the resource availability following the encoding of a high-frequency word at position *k*.

Once we have a validated neural index of resource availability, we can **use it to propose novel hypotheses regarding other long-term memory phenomena** that might be influenced by available cognitive resources during encoding. For example, successfully recalled items are associated with different neural patterns during encoding compared with those of unrecalled items. These differences are known as subsequent memory effects (SMEs) and they consistently appear across various brain regions, memory tasks, and imaging technologies (Kim, 2011; Paller et al., 1988; Paller & Wagner, 2002).

Though SMEs has been long established and robust, the interpretation of SMEs remains unclear (Halpern et al., 2021; Long et al., 2014; Paller & Wagner, 2002).

What are the underlying neural mechanisms during encoding that contribute to better recall performance? Multiple accounts have been provided in the past, linking SMEs to either endogenous factors such as attention and fatigue (Aly & Turk-Browne, 2016; Lohnas et al., 2020; Uncapher & Rugg, 2009) or external factors reflecting stimulus and task-related variables (Bainbridge et al., 2017; Craik & Lockhart, 1972; Fellner et al., 2013; Hanslmayr et al., 2009; Hanslmayr & Staudigl, 2014; Otten & Rugg, 2001a, 2001b; Sanquist et al., 1980). More recent efforts aim to further disentangle endogenous factors with external factors (Halpern et al., 2021; Kahana et al., 2018; Weidemann & Kahana, 2021).

In the current work, we show that the amount of available resources during encoding provides an alternative account for subsequent memory effects. Consistent with the SAC’s resource-depletion-and-recovery assumption, more available resources should relate to a more efficient encoding process and thus a better memory for an item. If the amount of available resources during encoding contributes to the observation of subsequent memory effects, we should observe a higher resource availability index for subsequently recalled items compared with subsequently unrecalled items. Moreover, according to the SAC’s resource-depletion-and-recovery assumption, as the resources recover gradually, the amount of resources available on one trial is correlated with the amount of resources at the preceding trial. Thus, we make a novel prediction about a potential sequential effect associated with the subsequent memory effect, which we termed as the “sequential subsequent memory effect”: if we split the subsequently recalled and subsequently unrecalled words and examine the neural activity when encoding the preceding item (in contrast to neural activity when encoding the given word as analyzed in standard “subsequent memory effect”), the resource availability index should be higher at encoding the preceding word among words that are subsequently recalled than words that are subsequently unrecalled. In other words, if the amount of resources available is high prior to the start of encoding an item, it is more likely that this item is subsequently recalled.

In the remaining paper, we provide the background of the SAC model from which we infer the amount of available resources during encoding and describe the procedure to obtain the neural index of resource availability from the EEG data. Then we validate the neural index of resource availability by comparing it against the inferred resources from the SAC model. Finally, we use the validated neural index as a tool to test a novel hypothesis that the subsequent memory effect is associated with the amount of available resources during encoding. Overall, these results provide converging support for the SAC’s resource assumption and a future tool for directly monitoring the amount of available resources from neural data.

## Method

### Participants

We analyzed the data from Experiment 4 of the Penn Electrophysiology of Encoding and Retrieval Study (PEERS) (Kahana et al., 2022). PEERS comprises 4 experiments and the datasets have been used in many other previous studies (Healey & Kahana, 2016; Healey et al., 2019; Lohnas et al., 2015). We excluded the participants for whom we could not find corresponding behavioral data and the final sample included 88 participants (age between 18-35) each with at least 20 completed sessions. The analyses in this study were not preregistered. We obtained pre-processed EEG data from Weidemann and Kahana (2021) and behavioral data from the lab that conducted the PEERS (Kahana et al., 2022; http://memory.psych.upenn.edu). We are grateful to the Kahana lab and to C. Weidemann for making their data available for reanalysis. The analysis code can be found at: https://rutgers.box.com/s/mfx50eea92e6giphgwck8405127p3vxu.

## Delayed free recall experiment

Participants performed a delayed free recall experiment with several experimental sessions consisting of 24 lists of 24 words. For each list, participants were asked to consecutively remember the word presented on the screen after which they performed a distractor task followed by a free recall task where they had 75 seconds to recall just-presented items in any order after 1200 - 1400 ms. At the beginning of each list, there was a 1500 ms delay before showing the first word. The presentation time for each word was 1600 ms, and the inter-stimulus interval between words randomly jittered between 800 - 1200 ms (uniform distribution). The distractor task lasted 24 seconds during which participants solved math problems in the form of A+B+C=? where A, B, and C are integers ranging from 1 to 9, and participants were given bonuses based on their speed and accuracy. Half of the lists were randomly chosen in each session (excluding the first list) where a pre-list distractor task was added. In the pre-list distractor task, participants performed the same math problem described above for 24 seconds before the presentation of the first word with an interval jittered between 800 - 1200 ms.

The experiment used a subset of 576 words drawn from an original word pool used in other PEERS experiments. For every participant, each session used these same 576 words but randomly assigned them to 24 lists. We obtained the word frequency of each word from the pre-generated word frequency data set from Semenov et al. (2015).

## EEG data collection and processing

Two systems, a 128 channel Biosemi Active and a 129 channel Geodesic Sensor (Electrical Geodesics, Inc.; EGI), were used in the experiment to record EEG data. Each participant used the same system for all their experimental sessions. The MNE package (Gramfort et al., 2013) and custom python codes were used to execute the raw EEG data.

A 0.1 Hz high-pass filter was applied to both systems and the EEG data were then re-referenced based on the average reference. The epoch of interest started from 800 ms before each word shown on the screen to the end of its presence. A 1200 ms buffer zone was applied to both sides of the epoch during wavelet transform after which a power spectrum analysis was conducted based on the logarithmically spaced 15 frequencies ranging from 2 to 200 Hz. A log-transform was applied to the power, the time was down-sampled to 50 Hz, and z-transformed powers were calculated across all trials within each session for each participant. As the goal of the study is to identify a single neural index of resources, and we do not have a strong hypothesis about which frequency bands at which electrodes are important, we averaged activity across electrodes in the superior region for our analysis.

These electrodes were used in previous studies (Paller et al., 1988; Weidemann & Kahana, 2021; Weidemann et al., 2009). For EGI system, the power at electrodes 4, 5, 12, 13, 19, 20, 24, 28, 29, 37, 42, 52, 53, 54, 60, 61, 78, 79, 85, 86, 87, 92, 93, 111, 112, 117, 118, and 124 were aggregated together, and for Biosemi system, electrodes A5, A6, A7, A18, A31, A32, B2, B3, B4, B18, B19, B31, B32, C2, C3, C4, C11, C12, C24, C25, D2, D3, D4, D12, D13, D16, D17, and D28 were aggregated together for subsequent analysis.

## Neural index of resource availability

The power spectrum patterns were averaged over a time interval of 0-1600 ms after each word presented. We defined this averaged power spectrum pattern as the key neural signature to derive the amount of available cognitive resources. According to the SAC model, participants have their full amount of available resources at the beginning of each list. Therefore, we used the power spectrum pattern at encoding the first word in lists as a template pattern for maximum resources. We reasoned that the more a power spectrum pattern is similar to the template pattern, the more resources are available. Since the power spectrum pattern is a vector of power values across frequencies, we calculated Pearson correlations between the template pattern and other power spectrum patterns as the measurement of the similarity. The Pearson correlation coefficient was then used as the resource availability index indicating how much cognitive resources are available. A larger divergence of a neural power spectrum pattern from the template corresponds to a lower index, indicating fewer cognitive resources available at that moment. The neural index varies between -1 and 1 with -1 indicating least resources and 1 indicating max resources. We compared the index in phenomena of interest across participants using mixed linear regressions and paired t-tests.

In every session of the experiment, half of the lists had a pre-list distractor task. We excluded those lists when calculating the neural template of maximum resources for the following reasons: first, completing the distractor task could cause additional depletion of cognitive resources prior to starting to learn a list; second, using lists without a preceding distractor task is a more typical free recall paradigm, and we hope this neural index can be applied as a tool for characterizing resources in more general cases. Because completing the distractor task could cause additional depletion of cognitive resources for early serial positions - we excluded the lists with a pre-list distractor task when comparing the neural pattern with respect to serial positions. Such distractor task does not influence the comparison of cognitive resources regarding word frequency and SMEs, therefore in those cases neural signatures were calculated across all lists to maximize statistical power.

## SAC model of resource availability

In the current work, we apply the SAC model of episodic memory to infer the amount of available resources during encoding, based on behavioral data during recall. The inferred resource availability will then be used to validate the neural index of resource availability, obtained from the neural signal during encoding. The current work does not represent novel theoretical model development of SAC per se - instead, it applies SAC to validate the resource-depletion-and-recovery assumption with neural data. That said, although it was not part of the original motivation behind this project, we were able to derive an analytic solution to SAC’s likelihood function, which has significant benefits for future model development.

We employed a simplified version of the SAC model (Popov & Reder, 2020) described below, that keeps the major assumptions about how the resources deplete and recover. The original SAC model is more complicated and involves assumptions about memory decay, contextual fan, and others; however, these assumptions are not relevant for the current work. Thus we decided to simplify the model in order to clearly demonstrate the resource-depletion-and-recovery assumption. This should allow for other computational models of memory to capture the same set of behavioral findings as long as the below assumptions of resources are incorporated into their models.

The model assumes that encoding information in LTM depletes a proportion of a limited resource pool (*W*_*max*_ = 1) and that the strength of the LTM trace is proportional to the amount of resources dedicated to it. In the model information is represented as a network of nodes that vary in strength between 0 and 1. Words have pre-existing semantic nodes whose strength *B*_*sem*_ is a continuous function of word frequency *f* measured per million words (fixed parameter values used by Popov and Reder, 2020):

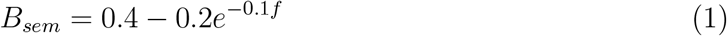

Based on this equation, we estimated that in the current experiment the strength of low-frequency words was 0.22, and the strength of high-frequency words was 0.40 (note that word frequency varied continuously, and the strength was also estimated continuously).

When encoding a word in a studied list, two processes occur. First, the activation of the semantic node is raised to 1, which depletes the following amount of resources:

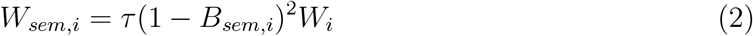

where *τ* is a scaling parameter and *W*_*i*_ is the amount of currently available resources at the begining of trial *i*. Thus, high-frequency words having higher base-level strength require less of the resource to be activated ((1 − 0.40)^2^*τ* = 0.360*τ*) relative to low-frequency words ((1 − 0.22)^2^ = 0.608*τ*). After the word is encoded, an episodic node is created which represents the binding between the word and the list context. Creating an episode node depletes a proportion *delta* of the remaining available resources, and its strength, *B*_*epi*_, is proportional to the amount of resources it consumed:

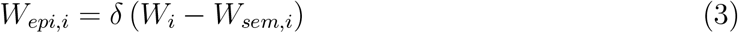

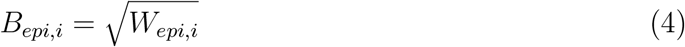

The resource pool recovers linearly over time *t*_*i*_ between item presentations at a rate of *r* per second until it reaches *W*_*max*_:

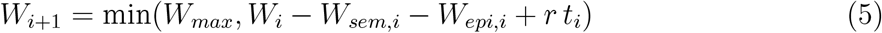

To transform memory strength, *B*_*epi*_ into recall probabilities, the model uses a standard signal detection rule. The model posits that successful recall of item i occurs probabilistically if the episode node’s strength is above the retrieval threshold:

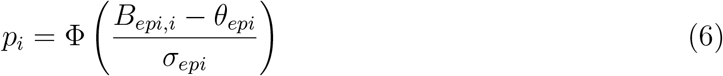

where *p*_*i*_ is the probability of succesful recall, *θ*_*epi*_ is the retrieval threshold, *σ*_*epi*_ is the standard deviation of normally distributed noise added to the episodic node’s strength, and Φ is the CDF of a standard normal distribution.

Finally, since individual recall attempts are coded as either successful (1) or unsuccessful (0), the likelihood of the response follows a Bernoulli distribution:

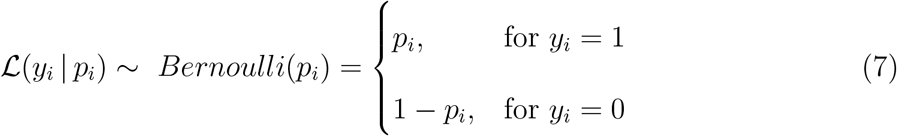

## Model fitting and simulation procedure

We first fitted the model to grouped behavioral recall data over serial position and word frequency of the preceding word (5 model parameters - *τ, δ, r, θ*_*epi*_ and *σ*_*epi*_). We only included serial positions 1-15 to estimate the model, because serial positions 16-24 showed recency effects. The reasons to exclude recency effects are two-fold. First, recency effects are not of interest in the current study, and including them or not does not affect how we compare resources obtained from the neural data versus SAC. Second, while it is possible to include a recency mechanism in SAC (Popov & Reder, 2020), using a simplified SAC without a recency mechanism critically allows us to derive the likelihood-based model that can be fit to individual subjects efficiently. We describe more details about individual model fitting in this section later. When fitting the model to behavioral patterns averaged across participants, five parameters were estimated via a grid search (*τ* = 0.2, *δ* = 0.26, *r* = 0.08, *θ*_*epi*_ = 0.367, *σ*_*epi*_ = 0.256) minimizing the root-mean-square error (RMSE) of the observed and predicted free recall probability. The obtained model gives us estimates of cognitive resources during the encoding of each word through *W*_*i*_ as found in Equation (6).

In addition to group-level model fitting, for each participant, we used maximum likelihood method to allow for efficient model parameter estimation at the trial level to obtain individual participant parameters. The full model likelihood function is derived in Appendix E. We used each individual model with the corresponding individual parameters to calculate the availability of cognitive resources at encoding the words with increasing serial positions. In the Results section, we report the group-level simulation on how resources change with the serial position and preceding word frequency. We also detail the results of individual-level simulation regarding serial positions in Appendix D and summarize it in the Results section.

The model so far only captures exogenous factors that affect resource depletion such as serial position and word frequency. To capture the effect of endogenous factors, in the Results section, we also report group-level simulations when there’s additional noise added to the resource depletion proportion on each trial (using the same set of model parameters previously fit to the group data). Specifically, we introduced normally distributed noise in the *delta* parameter on each trial *j*: *δ*_*j*_ ∼ 𝒩 (*δ*, 0.1), and an item is recalled if its episodic node’s strength is above the retrieval threshold *θ*_*epi*_.

## Results

Although the resource-depletion-and-recovery assumption of the SAC model has been supported by a series of behavioral findings (Kowialiewski et al., 2021; Mizrak & Oberauer, 2021; Popov et al., 2019; Popov & Reder, 2020; Popov et al., 2021), its neural underpinnings are yet to be found. We extracted a neural signature from EEG data and defined a corresponding neural index indicating the resource availability. To validate this resource availability index, we compared it with the resource availability inferred from the SAC model that captured the primacy serial position effect and the sequential effect of word frequency.

## Validation of resource availability index

First, we fitted the SAC model to the averaged recall data so as to see whether this simplified version based solely on the resource-depletion-and-recovery assumption could capture the behavioral pattern during recall. The model provided good fits to the primacy portion of the free recall data (Fig. 1; RMSE = 0.006, *R*^2^ = 0.989), with its estimated parameters consistent with previous SAC models (Popov & Reder, 2020). Specifically, as shown in the Fig. 1, the model, presented in the dash line, successfully captures the primacy serial position effect where the recall probability is the highest for the first presented word and decreases gradually with the serial position regardless of the word frequency of the preceding words. Meanwhile, at each serial position, the model captures the sequential effect of word frequency where the recall probability is higher when the word was preceded by a high-frequency word rather than a low-frequency word. As demonstrated in Appendix A, the model was also able to capture individual participant differences in the magnitude and duration of the primacy effect, as well as the word-frequency sequential effect when treated as a continuous rather than a binned binary variable.

**Fig. 1.**
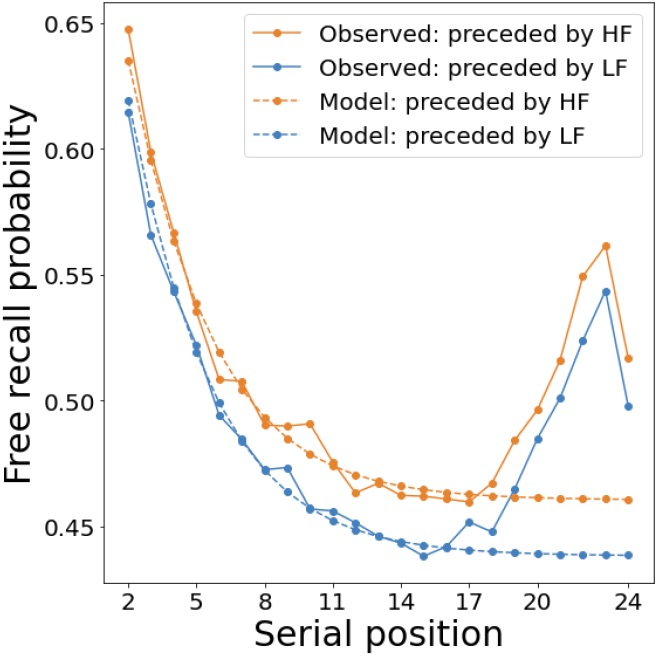
Primacy serial position effect (recall probability is at its highest at the beginning of the list) and sequential effect of word frequency (recall probability is higher when preceded by high-frequency words than low-frequency words), as captured by the SAC model. τ = 0.2, δ = 0.26, r = 0.08, θ_epi_ = 0.367, σ_epi_ = 0.256. Note that the model was only fit to serial positions 1-15 but we show the full curve for completeness and transparency.

With the SAC model fitted to the recall period of the behavioral data, we can infer the amount of available resources during encoding according to the SAC model and compared it with the neural index of resource availability (calculated separately from the encoding period of the neural data). Fig. 2a shows the amount of available resources at the beginning of encoding each serial position in SAC. At the beginning of the list resources are at their maximum, and they decrease gradually with each serial position, until they asymptote approximately after item 15. If the neural index indeed reflects the amount of available resources, it should follow the same pattern.

**Fig. 2.**
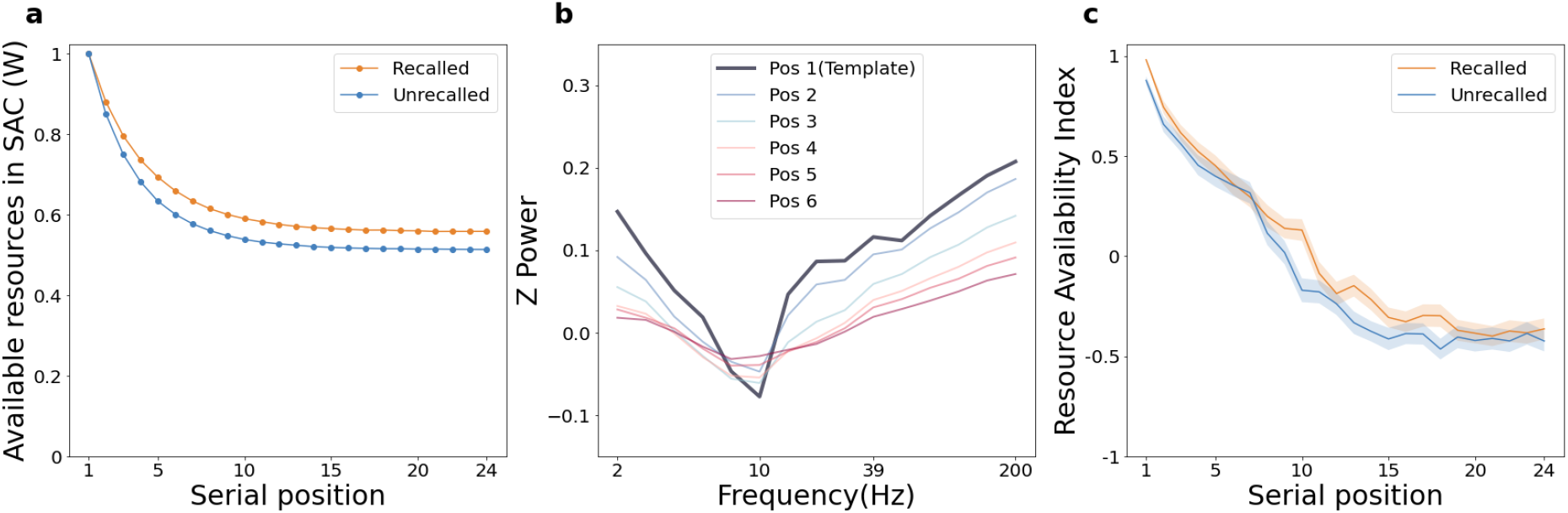
The effect of serial position on available cognitive resources during encoding. a. Available resources in the SAC model for different serial positions partitioned by whether words were subsequently recalled. b. The power spectrum pattern when encoding words from serial position 1 to 6 (in lists without a pre-list distractor task). c. Neural index of resource availability for different serial positions (in lists without a pre-list distractor task), partitioned by whether words were subsequently recalled. Note that: (1) figure a represents resources inferred from the model fitted to the recall data while figures b and c were measured from EEG data during the encoding period in the free recall task; (2) resources in the SAC model vary between 0 and 1, while the neural resource availability index varies between -1 and 1. These two notes also apply to corresponding sub-figures in Fig. 4, 5, and 6.

Fig. 2b shows the power spectrum pattern of neural patterns when encoding words at different serial positions (among lists without a pre-list distractor task). We assumed that the available cognitive resources are at their fullest at the beginning of each list and proposed the power spectrum pattern of encoding the first word in the list as a template pattern for maximum resources (Fig. 2b). We then measured the neural index of resource availability as how much the encoding patterns of other words are similar to this template pattern (Fig. 2c).

The template, i.e. serial position 1, has the lowest power value at the middle-frequency band and the highest value at the low- and high-frequency bands. As the serial position increases, generally, the value at the middle frequency band becomes larger and the value at the low- and high-frequency bands become lower (Fig. 2b). In other words, the neural signature diverges more from the template as the serial position increases, giving rise to decreased amount of resource availability as measured by our proposed neural index in Fig. 2c. Since recall performance highly correlates with serial positions, we plotted the pattern for recalled and unrecalled words separately. This pattern holds for both recalled and unrecalled words, demonstrating that the trend is not a trivial consequence of having higher accuracy at earlier serial positions. We fitted a linear mixed model to test the effect of serial positions on neural indexes. The effect of serial position is significant and negative in both recalled (beta = -0.06, 95% CI [-0.06, -0.05], *t*(2108) = -40.06, *p* < 0.001; Std. beta = -0.62, 95% CI [-0.65, -0.59]) and unrecalled condition (beta = -0.06, 95% CI [-0.06, -0.05], *t*(2108) = -38.83, *p <* 0.001; Std. beta = -0.59, 95% CI [-0.62, -0.56]). As the serial position number increases, the continuously decreasing index indicates fewer and fewer available cognitive resources during encoding, which matches the pattern of resource availability inferred from the SAC model in Fig. 2a.

Concerning the primacy effect, we note that SAC predicts an earlier convergence to its asymptote compared to the pattern demonstrated by neural data. This mismatch is possibly due to the simplified model setting which only focused on the resource-depletion-and-recovery mechanism. The model did not build any association between words, treated lists as independent instead of consecutive, excluded processes during retrieval, etc. All of these processes could potentially also deplete resources or bias the model parameter estimates (e.g. overestimate depletion rate). Per-participant level simulation also provides evidence that differences in the shape/steepness of the neural resource index can be explained by variations in resource parameters over participants.

Despite the early convergence, one might expect any natural drifting of neural signals leading to this decreasing function; however, this is not necessarily the case since neural drifting is not always monotonic across frequencies (see counter-examples in Appendix A).

If the model correctly captures the amount of available resources, fitting the model to individual participants should give us individual resource availability parameters that correlate with the magnitude of the neural resource index at the individual level. This would provide stronger evidence for the resource interpretation of the neural index. This is indeed what we found. For each participant, we calculated the average neural index collapsed over all trials, and we fit a linear regression model using the four model parameters as predictors (note that the *τ* parameter was fixed across participants, and that we had to exclude 32 participants because we could not uniquely identify a single set of parameters for them, see appendix D for more details). The regression model showed that the resource parameters of the model explain 20% of the variance in the overall neural index at the level of individual participants (see more details in Appendix D). Participants with larger *δ* parameter (i.e. the proportion of resources depleted by each item) have overall lower neural resource indices, shown by the negative correlation (*r*(55) = −0.758, *p <* 0.001) after controlling for the effect of other parameters (Fig. 3a). In Fig. 3b, we split participants into three groups based on their *δ* value and show their serial position curve separately. Likewise, groups with higher *δ* has lower neural indices at all serial positions. The effect of the recovery rate parameter is consistent with model prediction as well. After controlling for the effect of other parameters, we find a positive correlation between recovery rate and overall neural resource indices, *r*(55) = 0.75, *p <* 0.001 (Fig. 3c). After split into three groups based on recovery rate *r*, participants with higher recovery rate have higher neural indices at all serial positions (Fig. 3d). These per-participant level results significantly strengthen the support of the resource-based account.

**Fig. 3.**
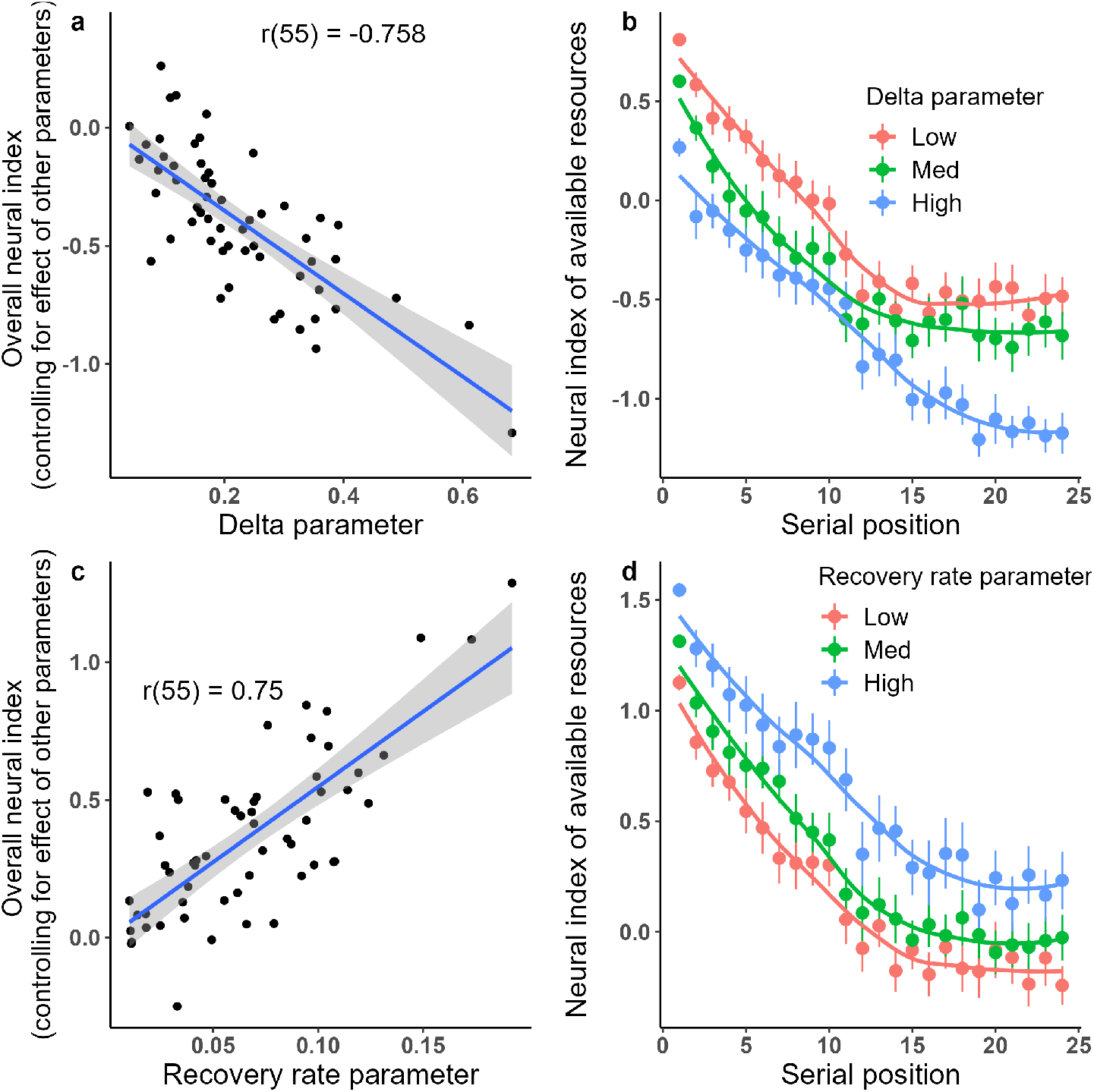
The effect of δ (the proportion of resources depleted by each item) and r (recovery rate) on available cognitive resources during encoding. a. Negative correlation between δ and overall neural index across participants. b. Participants were evenly split into 3 groups based on their δ parameters. Group with higher δ relates to an overall lower neural index. c. Positive correlation between r and overall neural index across participants. d. Participants were evenly split into 3 groups based on their r parameters. Group with higher recovery rate relates to an overall higher neural index.

After comparing available resources regarding serial positions, we next plot the amount of SAC inferred resources regarding the frequency of the preceding word. Fig. 4a shows the effect of word frequency at position k on the resource availability while encoding the immediately subsequent item (position k+1) or the item after that (position k+2).

**Fig. 4.**
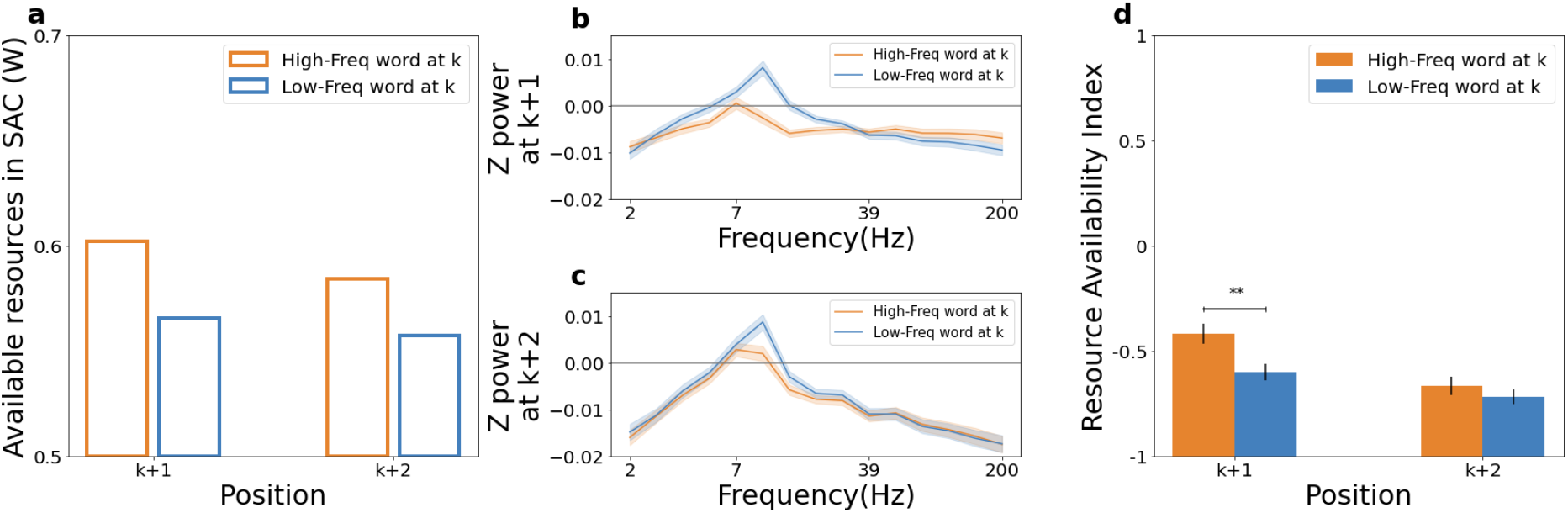
The effect of word frequency of the preceding word on available cognitive resources during encoding. a. Available resources in the SAC model during encoding words at positions k+1 and k+2 depending on the frequency of the word at position k. b. The power spectrum pattern when encoding words at position k+1 depending on the frequency of the word at position k. c. The power spectrum pattern when encoding words at position k+2 depending on the frequency of the word at position k. d. Neural index of resource availability during encoding words at positions k+1 and k+2 depending on the frequency of the word at position k. Asterisks indicate a significant difference between groups (**p < .01).

Specifically, studying low-frequency words at position k leaves fewer resources available when the next words are presented, and this effect is more pronounced during the encoding of words at position k+1 (*W*_*high*−*frequency*_ = 0.60, *W*_*low*−*frequency*_ = 0.57) than during the encoding of words at position k+2 (*W*_*high*−*frequency*_ = 0.58, *W*_*low*−*frequency*_ = 0.56), as the latter has more time to recover the pool of cognitive resources. If the neural index we identified represents resource availability as we hypothesized, then it should show similar patterns as a function of word frequency as the SAC model’s inferred resources based on recall performance (Fig. 4a).

Fig. 4b and c show the power spectrum patterns when encoding words at position k+1 and k+2. We split trials into 5 bins with equal number of words in each based on their word frequency at position k (k range from 1 to 22) and compared the corresponding neural signatures in bins of the highest and lowest word frequency. In both Fig. 4b and Fig. 4c, neural patterns relate to high-frequency words at position k are more similar to the template representing maximum resources (see Fig. 2b). Because the neural index of resource availability is defined as the similarity to the template, this means that there are more resources available when word encoding is preceded by a high-frequency word (Fig.4d). To formally test whether higher neural index is related to encoding words following high-frequency rather than low-frequency words and whether this effect is more pronounced at position k+1 than k+2, we performed a two-way repeated ANOVA on the neural index. The result shows a significant effect of interaction between word frequencies and serial positions (F(87)=3.97, p<0.05). In comparison to encoding low-frequency words (M=-0.60, SE=0.04), encoding high-frequency words (M=-0.43, SE=0.05) at position k is associated with a significant higher resource index value (t(87)=2.81, p=0.006, d=0.42) at position k+1. Encoding words of different word frequencies does not have a significant effect on the neural index at the k+2 position. Altogether, high-frequency words at position k are related to more available cognitive resources at position k+1 and a smaller, even though non-significant, facilitation at position k+2, which matches the pattern of resource availability inferred from the SAC model in Fig. 4a. Comparison between the low-frequency bin and the high-frequency bin was chosen because of the ease of visual demonstration. In the previous analysis we split word frequency into bins, because it makes it easier to visualize the trend. However, word-frequency is a continuous variable, and in principle the model takes into account each specific item’s frequency, rather than its bin. In Appendix B, we applied mixed linear models and confirmed the significant effect of word frequency in the analysis of both finer frequency bins and specific word frequencies.

Both the evidence from the primacy serial position effect and the sequential effect of word frequency validated that our proposed neural index of resource availability reflects cognitive resource availability. This conclusion is further supported by the correlation between resource parameters for individual participants, and the magnitude of their individual neural index. Following the validation, we are interested in whether the neural index can serve as a tool to examine long-term memory phenomena that might be influenced by available cognitive resources during encoding.

## An alternative account for subsequent memory effects

Subsequent memory effects (SMEs) refer to the observation that neural patterns of encoding subsequently recalled items are different with that of encoding subsequently unrecalled items. Multiple accounts have been proposed in the literature, associating SMEs with attention, fatigue and properties of stimuli (Aly & Turk-Browne, 2016; Craik & Lockhart, 1972; Halpern et al., 2021; Hanslmayr & Staudigl, 2014; Lohnas et al., 2020; Otten & Rugg, 2001b; Uncapher & Rugg, 2009; Weidemann & Kahana, 2021). We propose that observed SMEs are associated with different amount of available cognitive resources during encoding.

We first reproduced in Fig. 5a the subsequent memory effects commonly observed in the literature, where the neural patterns for subsequently recalled words are different than those for subsequently unrecalled words. Consistent with prior studies (Gruber et al., 2004; Hanslmayr et al., 2012; Sederberg et al., 2006; Sederberg et al., 2003; Summerfield & Mangels, 2006), Fig. 5a demonstrates that the power spectrum pattern associated with subsequently recalled words has a lower value at the middle-frequency band and higher values at the low-frequency and high-frequency bands. Unique to our current analysis, however, is our ability to interpret these patterns as how similar they are to the template that represents maximum resources, i.e. there is a smaller divergence from the first encoded word template among the recalled words compared to unrecalled words, indicating that there are more resources available when encoding words that are subsequently recalled.

**Fig. 5.**
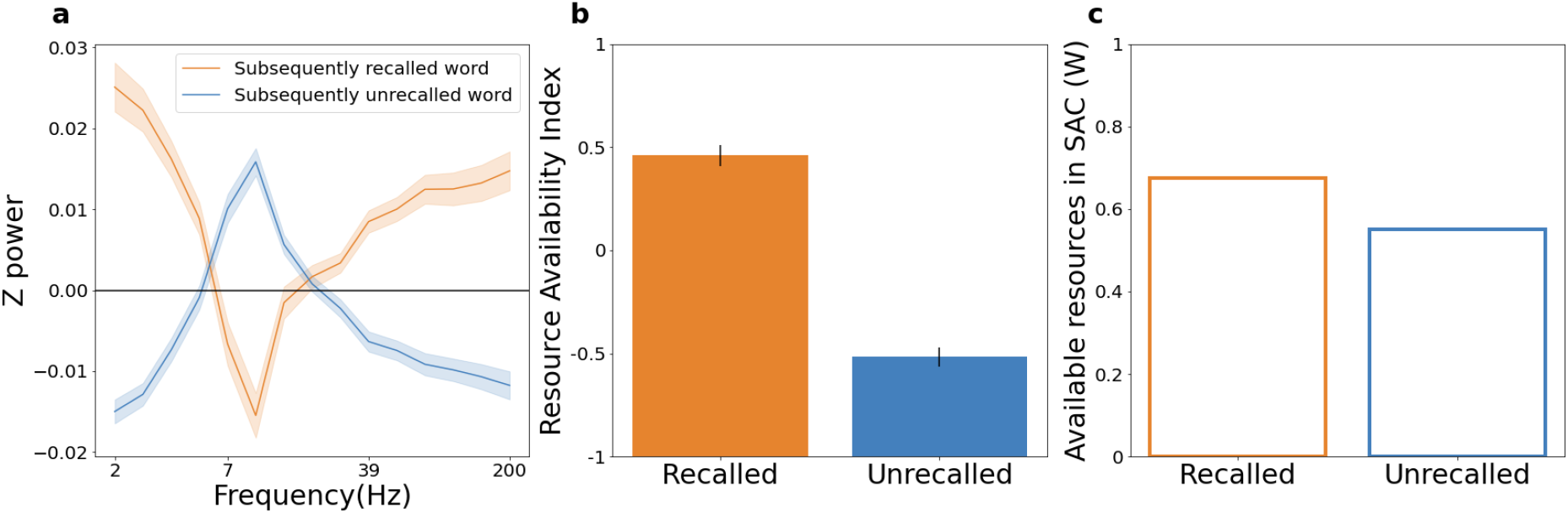
Explaining subsequent memory effects by cognitive resources available during encoding. a. Different power spectrum patterns for encoding subsequently recalled versus unrecalled items (i.e. subsequent memory effects). b. The neural index of resource availability of subsequently recalled versus unrecalled words. c. Available resources in the SAC model during the encoding of subsequently recalled versus unrecalled words.

This effect is more clearly seen in Fig. 5b, where there is a significant higher neural index of resource availability (t(87)=9.93, p<0.001, d=2.07) at encoding subsequently recalled words (M=0.76, SE=0.09) than subsequently unrecalled words (M=-0.86, SE=0.08). One may wonder if this observation provides any additional unique evidence to support the resource account for the neural index, since we have already shown that earlier serial positions in the list are associated with higher neural index, and that recalled words are more likely to come from earlier serial positions. This is already clear from Fig. 2c, which dissociates serial position from recall status; to statistically validate the claim that recalled words are associated with higher neural index than unrecalled words, even after controlling for serial positions, we ran a mixed linear model with random intercept and slope to control the effect of serial position. The effect of the recall (recalled word was encoded as 1 otherwise 0) is statistically significant and positive (beta = 0.08, 95% CI [0.06, 0.11], *t*(4217) = 6.71, *p <* .001; Std. beta = 0.06, 95% CI [0.05, 0.08]). Again, the result indicates that compared to unrecalled words, recalled words are associated with higher neural index.

This result supports our hypothesis that the differential neural patterns at encoding for subsequently recalled versus unrecalled words could be a result of participants having different cognitive resources available during encoding. We also simulated available resources during encoding of subsequently recalled or subsequently unrecalled words according to the same SAC model we fitted before (Fig. 5c). Fig. 5c shows that the available resources in SAC are higher during encoding subsequently recalled words compared to subsequently unrecalled words (*W*_*recalled*_ = 0.67, *W*_*unrecalled*_ = 0.55), consistent with the neural index pattern.

To further support our hypothesis that the SMEs could be a result of participants having different cognitive resources available during encoding, we made a novel prediction of a phenomenon that is closely related to the SMEs, which we call the “sequential subsequent memory effects” - subsequent better memory for item *X*_*k*+1_ is associated with different neural patterns when encoding *X*_*k*_ (instead of *X*_*k*+1_ as in standard SMEs; k and k+1 are the positions of the item). This is shown in Fig. 6a where the power spectrum pattern for encoding words at position k is different for subsequent recalled versus unrecalled items encoded at position k+1; this holds either when items encoded at position k were subsequently recalled (Fig. 6a) or unrecalled (Fig. 6b). Importantly, we predict that better memory for item *X*_*k*+1_ is associated with more resources during the encoding of the preceding item *X*_*k*_, as the available resources during encoding words at position k only change gradually and could carry over to position k+1.

**Fig. 6.**
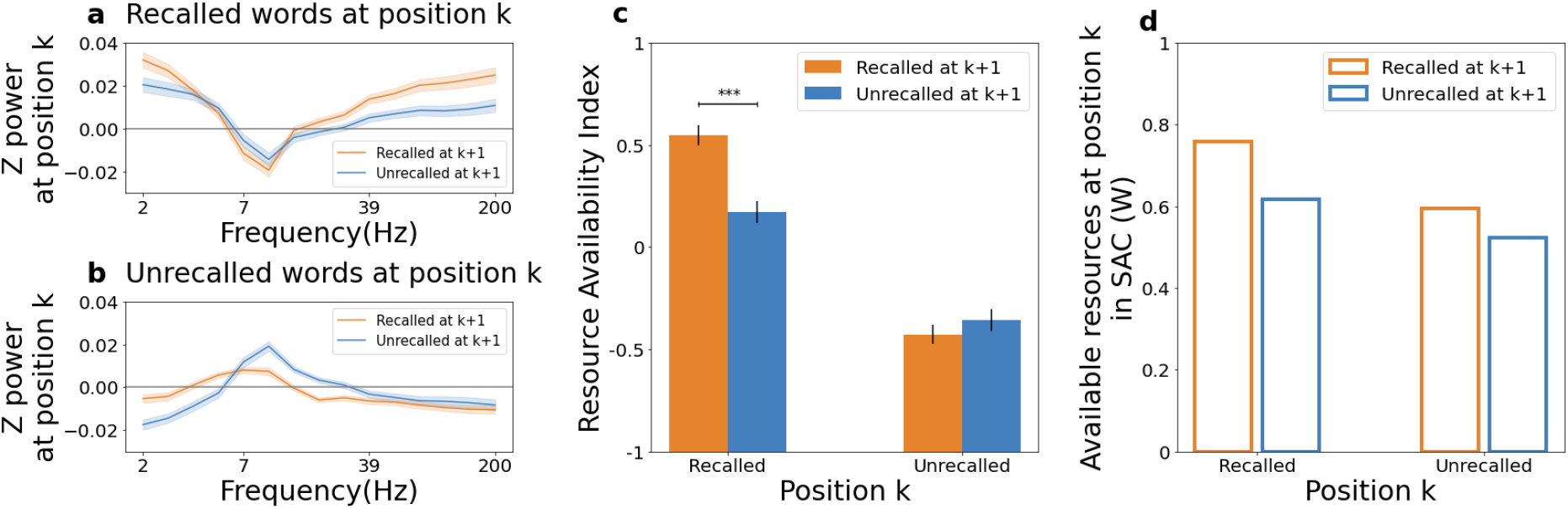
Explaining sequential subsequent memory effect by cognitive resources available during encoding. a. Different power spectrum patterns when encoding recalled words at position k for subsequently recalled versus unrecalled items encoded at position k+1, i.e. sequential subsequent memory effects, conditioned on when items at position k were subsequently recalled. b. Different power spectrum patterns when encoding unrecalled words at position k for subsequently recalled versus unrecalled items encoded at position k+1, i.e. sequential subsequent memory effects, conditioned on when items at position k were subsequently unrecalled. c. The neural index of resource availability when encoding items at position k, conditioned on whether items at position k and k+1 were subsequently recalled or unrecalled. Asterisks indicate a significant difference between groups (***p < .001). d. Available resources in the SAC model when encoding items at position k, conditioned on whether items at position k and k+1 were subsequently recalled or unrecalled.

We tested this prediction by plotting the neural index of resource availability during encoding of *X*_*k*_ given whether *X*_*k*+1_ were subsequently recalled or not (Fig. 6c). When the word *X*_*k*_ was recalled, more resources are related to successful recall of the word *X*_*k*+1_ (M=0.568, SE=0.049), with a significantly higher index value (t(87)=6.367, p<0.001, d=0.774) in comparison to where the word *X*_*k*+1_ was not recalled (M=0.192, SE=0.055).

When the word *X*_*k*_ was unrecalled, the resource availability indexes are not significantly different from each other (Fig. 6c) - possibly due to an already nearly fully depleted resources indicated by the preceding state where participants had already failed to recall. Altogether, these neural results show that more cognitive resources available at encoding preceding words relate to better memory of current words, indicating that the amount of resources available at any point is associated with not only the memory of the item encoded at that point, but also the memory of items encoded in the near future. Even when there are enough resources at time k to encode item *X*_*k*_ successfully, having even more available resources would leave enough for encoding the subsequent item *X*_*k*+1_.

To test whether the SAC model also produces these sequential SME effects, we simulated how the resources change in such condition according to the already fitted SAC model (Fig. 6d). The simulation result has a close correspondence with the neural index (Fig. 6a). Specifically, more resources are available during encoding *X*_*k*_ when *X*_*k*+1_ was recalled rather than unrecalled and such effect is larger when *X*_*k*_ is recalled (*W*_*recalled*_ = 0.76, *W*_*unrecalled*_ = 0.62) compared to when *X*_*k*_ is unrecalled (*W*_*recalled*_ = 0.60, *W*_*unrecalled*_ = 0.52). The resource availability inferred from the SAC model (Fig. 6d) corresponds to that measured by our neural index during encoding (Fig. 6c), although the model does predict a small difference even when item *X*_*k*_ is unrecalled.

While the sequential SME effects are consistent with the SAC model, there are two possible confounding factors: serial position and contiguity effect. If item *X*_*k*_ and *X*_*k*+1_ were both recalled, they are more likely to be words presented earlier than later. Since earlier serial positions are related to higher neural index values compared to later serial positions, the higher index value shown in the sequential subsequent memory effect may be solely due to serial positions at which words were presented. To control the effect of serial position, we ran a mixed linear model with random intercept and slope. The dependent variable is the neural index at item *X*_*k*_. The predictors are whether item *X*_*k*_ was recalled, whether item *X*_*k*+1_ was recalled, and the serial position. The result shows a positive significant effect of whether k+1 was recalled (beta = 0.02, 95% CI [1.99e-03, 0.04], *t*(8088) = 2.17, *p* = 0.030; Std. beta = 0.02, 95% CI [1.57e-03, 0.03]), indicates that the sequential subsequent memory effect is not solely due to the serial position.

The second factor is the contiguity effect during recall. The contiguity effect indicates that when an item *X*_*k*_ is retrieved during the recall phase then the next retrieved item is likely to be *X*_*k*+1_ (Kahana, 1996). Thus the existing difference which is based on the comparison between item *X*_*k*_ and *X*_*k*+1_ might be caused by something related to retrieval not resources during encoding. We conducted additional analysis while controlling for the contiguity effect (see Appendix C), which did not change our conclusions.

Overall, the novel “sequential subsequent memory effects” that we observed and their corresponding neural index of resource availability serve as further evidence that the amount of available cognitive resources provides an alternative account for the subsequent memory effects.

## General Discussion

Humans possess a limited amount of cognitive resources to process various cognitive operations (Cary & Reder, 2003; Diana & Reder, 2005; Popov & Reder, 2020; Reder et al., 2000; Reder et al., 2007); however, the neural substrate of the resources is not known. In the current study, we identified a neural index during word encoding in a free recall paradigm which can be interpreted as reflecting the amount of cognitive resources available for forming new memory traces. We validated this neural index of resource availability (measured from the neural data during encoding period) by demonstrating its alignment with inferred resources from the SAC model (obtained from the behavioral data during the recall period): resource availability in both SAC and neural index decreases with serial positions, and is higher following encoding a high-frequency word than a low-frequency word. Finally, by fitting the model to individual participants, we found that participants with higher resource-depletion rates or slower resource-recovery rates, as estimated by the model, also had a lower neural index. Combined, these effects provide substantial support for our resource availability interpretation of the neural index identified in the current paper.

The advantages of separately obtaining the neural resource availability and the model-based resource availability before evaluating their alignment is two folds. First, it is challenging to directly interpret what changes in the neural index represent; alignment between the neural index and the well-specified property of cognitive resource in SAC allows us to interpret the obtained neural index as the amount of available resources. This alignment further allows us to explain the long-established subsequent memory effects (SMEs) in terms of the amount of cognitive resources available during encoding, as we observed higher neural index for subsequently recalled items compared with that for subsequently unrecalled items. Second, the discovery of a neural index that aligns with how cognitive resources changes in SAC provides converging support to the proposed limited resources in the SAC model, especially as the neural index of resource availability and SAC’s resource availability are derived from separate portions of data. We now turn to the broader implications of these results.

## Distinguishing from alternative interpretations of the neural index

One important feature that distinguishes the SAC’s resource explanation from other constructs like attention is its well-specified mathematical property, rather than a vague appeal that “more attention is devoted to items that are subsequently recalled”. This is a nontrivial difference - the precise specification of resources allows us to test a variety of phenomena and derive numerical predictions as we have already shown. For these predictions, the SAC’s resource assumption successfully explained all but other constructs could only account for a subset.

For example, one may wonder if the identified neural index reflects the rhythmic waxing and waning of attention. However, rhythmic changes in attention predict that there should be autocorrelation in attention from trial to trial, but not that attention should always decrease over time and be reinstated at the beginning of the list. One could of course impose a monotonic decrease function of attention, as in the Context Retrieval and Maintenance model (CMR; Polyn et al.,2009), but there is no principled reason behind this other than a convenient mathematical function to account for the primacy effect. In contrast, the SAC resource depletion mechanisms naturally give rise to this monotonic function, because each item depletes a fixed proportion of the remaining available resources, which, together with the gradual recovery of resources leads to initial steeper depletion, then eventually to an asymptote. Thus the SAC model derives this monotonic function from a mechanism, instead of just stipulating its existence. Additionally, there are other issues with the attentional fluctuations account. Attentional fluctuations cannot explain the sequential effect of word frequency in the manuscript. Low-frequency words attract more attention relative to high-frequency words (Diana & Reder, 2005; Malmberg & Nelson, 2003; Popov & Reder, 2020). If the neural index represents rhythmic waxing and waning of attention, then the neural index should be higher on items that follow low-frequency instead of high-frequency items, because attention is still in a heightened state. We found exactly the opposite, which is consistent with the SAC account, because in SAC once more resources are depleted by the low-frequency words, fewer resources are available for processing subsequent items.

Another alternative is the context change account, as context drifts gradually from encoding one item to the next (Howard & Kahana, 2002). One can interpret the neural index as reflecting the amount of context change since the beginning of the list. As more surprising events may lead to greater context change and low-frequency words are generally more surprising, one might explain the lower neural index after encoding low-frequency words than high-frequency words as having a larger context change. However, the same context change account cannot explain why this effect is smaller two positions, instead of one position, after encoding a low/high-frequency word; in contrast, the resource account is associated with a mechanism where depleted resources can recover over time, accounting for a smaller effect for words presented two positions after high-frequency and low-frequency words. Additionally, the context change account does not explain why the neural index varies as a function of subsequent memory. We found that words that are subsequently forgotten have a lower neural index during encoding relative to words that are subsequently remembered. If the neural index represents the amount of context change during encoding relative to the start of the list, the difference between indexes during encoding indicates that subsequently forgotten words relate to a larger context change. It is possible that some words naturally induce greater context change, but no evidence says they should then be worse remembered. If anything, we might expect the opposite, since encoding items in more distinct contexts should make them easier to retrieve due to their distinctiveness (Siefke et al., 2019).

Perhaps most importantly, we believe that the per-participant simulations of the SAC model and their correlations with the neural index provide further support for the model (see Appendix D). For example, participants with faster resource recovery rates had higher overall neural resource indices - this effect cannot be explained by endogenous factors such as contextual drift or external factors such as serial position. While alternative explanations might account for certain subsets of the data, taken together, only the resource-based account account for the entire spectrum of results.

## A model-based approach to identify neural substrates

Typical cognitive neuroscience approaches focus on applying statistical models to determine which aspect of the neural data are associated with certain behavioral outcomes or experimental manipulations (Turner et al., 2017). For example, by contrasting neural patterns during a “2-back” working memory task and a control task that requires only perceptual and response processes, one can identify a set of brain regions responsible for working memory maintenance (see a review at Smith and Jonides, 1998). Similarly, SMEs have been studied by contrasting neural patterns during encoding for later recalled items against those of unrecall items: Using EEG and intracranial EEG, different brain sites were shown to be related with the successful memory in gamma and theta oscillatory activities either controlling the serial position (Sederberg et al., 2006) or not (Sederberg et al., 2003). Likewise, Sederberg et al. (2007) found that the same pattern of gamma activity observed across various brain regions during encoding reappeared later during retrieval. These studies uncover useful information about the brain regions and oscillatory activities involved in SMEs, however, it is challenging to interpret why these regions are involved and the exact mechanism of that involvement (Palmeri et al., 2017; Turner et al., 2017). What is more, sometimes it is unclear what behavior outcomes or experimental manipulations one should contrast the brain activity with. For example, to apply a similar approach, our current study could design an experiment with two conditions - corresponding to “high” and “low” demand of cognitive resources - and compare neural patterns between the conditions. However, cognitive resource is a latent cognitive construct that we cannot easily postulate its connection with behavioral outcomes or experimental conditions. What could be designated as “high cognitive resources” and “low cognitive resources” conditions could involve other cognitive processes that result in a complex relationship between the factors we are interested in in the behavioral outcomes to predict (see analyses that demonstrated such complication; Price et al., 1996; Zhang et al., 2018).

To tackle this problem, we adopt a model-based approach by comparing our proposed neural index of cognitive resources with a latent quantity of cognitive resources in a computational model of memory, i.e. SAC. In contrast to cognitive neuroscience approaches, models of cognition have the advantage to characterize latent cognitive processes in precise mathematical terms. The amount of available cognitive resources in SAC during encoding were decided so that the model captures behavioral findings in free recall that are associated with the recovery and depletion of cognitive resources.

Additionally, while models of cognition typically rely on comparing their fit to behavioral findings to test alternative theories (Roberts & Pashler, 2000), being able to identify the neural substrate of a cognitive process in the model can provide further support for the theory that specifies the cognitive process (Anderson et al., 2016; Huber et al., 2008; Jacob & Huber, 2020; Nunez et al., 2019; Nunez et al., 2017; Padoa-Schioppa, 2011; Zhang et al., 2017). Importantly, instead of finding the neural index as the highest correlated neural signal with predictions from the SAC, we derived it independently from the encoding phase of the neural data before demonstrating its alignment with resources inferred by the SAC from the recall phase of the behavioral data. Since two different sources of data (encoding and recall) are used to identify the neural resource availability index and the model’s latent resource availability quantity, this approach provides strong convergent evidence for the model’s resource assumptions (see a recent discussion of evaluative standards for models of memory proposed by Popov, 2023).

## Reconcile accounts for subsequent memory effect: endogenous or exogenous factors

To better understand the neural mechanisms during encoding that contribute to SMEs, it is helpful to distinguish endogenous factors that reflect endogenously varying brain states with external factors reflecting stimulus and task-related variables (Weidemann & Kahana, 2021). Weidemann and Kahana (2021) identified the effect of serial position on neural activity, i.e., neural activity at early serial positions is similar to those of subsequently recalled items and that at later serial positions resembles that of subsequently unrecalled items. These were taken as evidence that the standard subsequent memory analysis may be misleading (differences in neural activity between subsequently recalled and subsequently unrecalled items might reflect processes controlled by exogenous factors such as serial positions) and that one needs to analyze SMEs in a way that isolates exogenous factors from endogenous factors (Weidemann & Kahana, 2021). In a novel analysis approach of SMEs, Halpern et al. (2021) also emphasized the importance of adjusting for exogenous factors such as serial positions and item effects. However, our current work demonstrates that endogenous and exogenous factors may not be easily separated. This is because exogenous factors such as serial positions can directly covary with an endogenous factor such as fluctuations in the amount of cognitive resources available: according to the SAC’s resource-depletion-and-recovery assumption, when resource depletion is faster than resource recovery, as the list moves forward, there are gradually less and less cognitive resources available. This also simultaneously explains why neural activity at early serial positions is more similar to those of subsequently recalled items than unrecalled items, since both the early serial position items and the subsequently recalled items are associated with more resources.

It is important to note that SMEs might represent a family of effects, rather than a single unified signal, and that multiple processes might be contributing to observed ERP patterns during the encoding of recalled vs unrecalled items (Van Petten & Senkfor, 1996). Our findings suggest that at least part of SMEs signal can be attributed not to encoding processes per se, but to the “readiness to encode”, consistent with prior findings that EEG activity before an item is presented can predict subsequent memory of that item (Noh et al., 2014). The novel sequential SME effect we uncovered based on the SAC’s predictions is particularly interesting in this regard. We showed that even when item *X*_*k*_ is successfully recalled, the neural patterns during its encoding differ depending on whether the subsequent item, *X*_*k*+1_, was recalled. If SMEs represented processes entirely related to the encoding of the current item, such as its memorability, salience, or the strength of the resulting memory trace, then there is no reason to expect a difference based on the recall probability of the subsequent item.

## Limitations and future work

The SAC’s resource-depletion-and-recovery assumption is agnostic about the nature of the cognitive resources - in it, resources are a latent quantity that gets assigned as “fuel” to each memory operation. In our current work, we demonstrated that this is more than just an abstract mathematical quantity but that it corresponds to concrete neural substrates. While the current results take us one step closer to understanding the neural underpinnings of cognitive resources, it is important to note that as of yet they do not unambiguously tell us what the nature of these resources is. Are these cognitive resources the same resources that underlie one’s limited capacity during working memory tasks (Anderson et al., 1996; Ma et al., 2014; Oberauer et al., 2016)? Are these cognitive resources related to the inner resources drawn during self-regulation in the influential “ego-depletion” account (Bratslavsky et al., 1998) (though recently called into question through multiple meta-analyses from Carter et al.,2015; Hagger et al.,2016; Randles et al.,2017)? Both are excellent venues for future research.

Besides exploring the nature of resources, one might also be interested in a more comprehensive analysis of neural patterns. In the current study, we used a simplified approach to identify a neural resource availability index by averaging across electrodes and time. While this achieved the purpose of identifying a relevant neural index, analyses such as brain activity localization or using a more fine-grained temporal resolution could be helpful in future work.

In addition to providing the first neural support of the SAC’s resource-depletion-and-recovery assumption, the current results also provide a potential methodological tool with which to study resource availability in the future. One drawback of behavioral studies is that differences in resource availability during encoding can only be inferred based on differences in subsequent recall performance. In the past this has significantly limited the questions we can ask about resource depletion. For example, for modeling simplicity, Popov and Reder (2020) assumed that resource depletion during encoding occurs instantaneously with the word presentation and that afterwards the resources recover linearly over time. As they noted in the conclusion section, these choices allowed the model to fit the data, but were otherwise arbitrary. With the currently proposed resource availability index, we could in the future potentially directly measure the time dynamics of resource depletion and recovery during encoding.

## Conclusion

While there have been numerous behavioral findings that support an account of limited resources during memory encoding, neural underpinnings of the resources are unknown. Our work fills this gap by identifying a neural index of resource availability extracted from EEG data. We validated this neural index by showing its consistency with inferred resource availability from the SAC model that was able to capture the primacy serial position effect and the sequential effect of word frequency. Further, we used this neural index as a tool for monitoring resource availability and showed it was able to provide an alternative account for the long-established subsequent memory effects. Together, these results provided converging support for the proposed limited resources in the SAC model and a future tool for directly monitoring the resource availability from neural data.

## Appendix A Additional analysis considering alternative accounts of the neural index

As reported in the main text, the neural index continuously decreases as the serial position increases, eventually reaching an asymptote. One might argue that the neural signal drifts away from the original point as time passes by and defining the power spectrum pattern at encoding the first word in lists as the template pattern seems to necessarily produce such a decreasing function. If this were the case, it would significantly undermine the support this curve provides for the resource model. However, this is not the case since neural drifting is not always monotonic. For example, the alpha band power values follow a non-monotonic function where the power values first decrease and then increase (Fig. A1). Fig 2 in Sederberg et al. (2006) similarly demonstrated this fact - not all frequency bands give a monotonic trend with serial positions. The neural index was measured as Pearson correlation coefficients between template pattern and power spectrum pattern at encoding other words. Since the power value function at different frequency bands does not monotonically increase or decrease, the template we currently use should not necessarily produce a continuous decreasing trend in neural index.

**Fig. A1.**
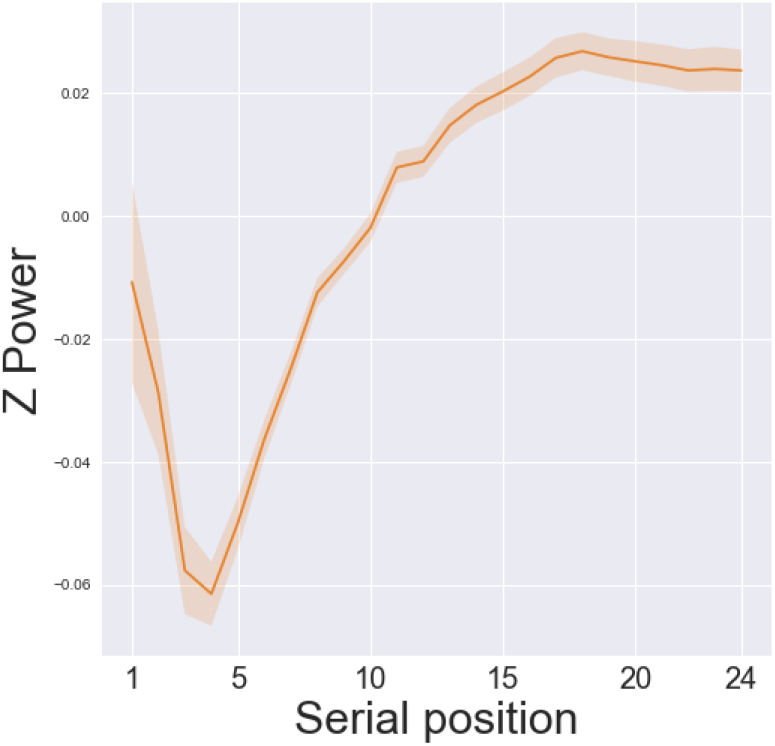
The effect of serial position on the alpha band power value during encoding. Values first decrease and then increase.

## Appendix B Additional analysis of the neural index using finer word frequency bins

We performed additional neural index analysis for finer word frequency bins and also for specific word frequencies separately. We partitioned words into 5 and 20 bins with respect to their word frequency where each bin had the same number of words. For example, in the 5-bin case, the first bin contained the first 20% of the lowest frequent words. We then performed random intercept mixed linear models using the neural index at item *X*_*k*+1_ as the dependent variable and word frequency of *X*_*k*_ as the predictor. The results (Fig. A2) showed a significant positive effect of word frequency in both 5-bin (beta = 0.04, CI = [0.01, 0.06], t(436) = 2.81, p= 0.005; Std. beta = 0.12, 95% CI [0.04, 0.20]) and 20-bin (beta = 5.76e-03, CI = [1.96e-03, 9.55e-03], t(1756) = 2.97, p= 0.003; Std. beta = 0.07, 95% CI [0.02, 0.11]). Similar to the previous behavioral study (Popov & Reder, 2020), the effect of frequency is smaller in finer frequency bins as data is more noisy.

We further conducted analysis on word frequency values without using frequency bins. Before fitted to the mixed linear model, the word frequency was log-transformed. The result shows that the effect of frequency is statistically significant and positive (beta = 4.67e-03, 95% CI [2.16e-03, 7.18e-03], *t* (50684) = 3.64, *p <* .001; Std. beta = 0.02, 95% CI [7.45e-03, 0.02]).

**Fig.A2.**
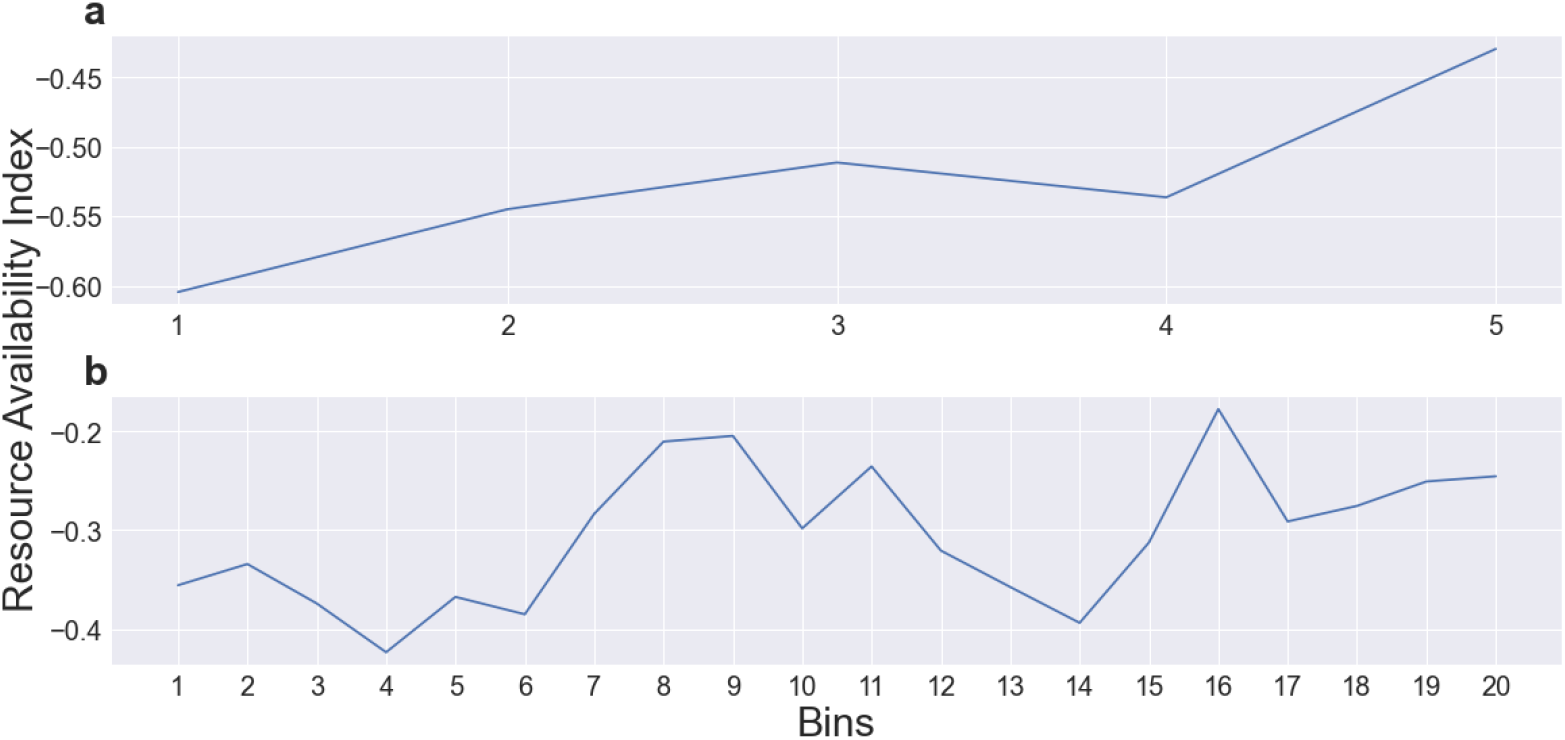
Neural index of resource availability during encoding words at positions k+1 increases as a function of word frequency at position k, partitioned into 5 bins (a) or 20 bins (b).

## Appendix C Additional analysis considering alternative interpretation of the sequential subsequent memory effect

In the sequential subsequent memory effect, the results showed that more available resources at recalled item *X*_*k*_ is associated with better memory at item *X*_*k*+1_. However, one potential confound in this effect is the contiguity effect in recall probability between nearby items. Specifically, the contiguity effect indicates that when item *X*_*k*_ is retrieved then the next retrieved item is likely to be *X*_*k*+1_ (*k* and *k* + 1 is the serial position during encoding; Kahana, 1996). Therefore the existing difference on the neural index based on the comparison between encoding item *X*_*k*_ and *X*_*k*+1_ might be influenced by factors related to the retrieval phase. To rule out this interpretation of the identified neural index, we separated the case where both item *X*_*k*_ and *X*_*k*+1_ were recalled into “far” and “near” conditions based on the recall phase. The “near” condition included item *X*_*k*_ and item *X*_*k*+1_ that were recalled adjacently and the “far” condition included words that were recalled far apart, i.e., all non-adjacent recalls.

We repeated the analysis. Compared to Fig. 5, Fig. A3a shows additional results where the power spectrum pattern for encoding words at *X*_*k*_ is different in the “far” condition and “near” condition. The neural index value is shown in Fig. A3c. In both “far” and “near” conditions, more resources are related to successful recall of the next word *X*_*k*+1_ (“near”: M=0.56, SE=0.05; “far”: M=0.44, SE=0.05), with significantly higher index values (“near”: t(87)=5.83, p*<*0.001, d=0.76; “far”: t(87)=4.34, p*<*0.001, d=0.50) compared with where the word *X*_*k*+1_ was not recalled (M=0.19, SE=0.05). The “near” condition has a significantly higher index than the “far” condition (t(87)=2.11, p=0.04, d=0.25).

Since the neural index is higher when *X*_*k*+1_ is recalled relative to unrecalled, regardless of the recall order, the output order is not a confound to the sequential subsequent memory effect. Moreover, the difference between the neural index during the encoding of items that were recalled near or far apart later can be interpreted in the SAC framework. According to the latest versions of SAC, when items are encoded, they are bound to their context, which includes information about other items. The strength of these item-context links determines the strength of the contiguity effects (Popov & Reder, 2020). In turn, the amount of resources available at encoding determines the strength of those item-context links. Therefore, a prediction of the verbal model (although not implemented computationally yet), would be that when there are more resources available during the encoding of two neighboring items, these items would be more likely to be recalled together later, which is consistent with this additional analysis.

**Fig.A3.**
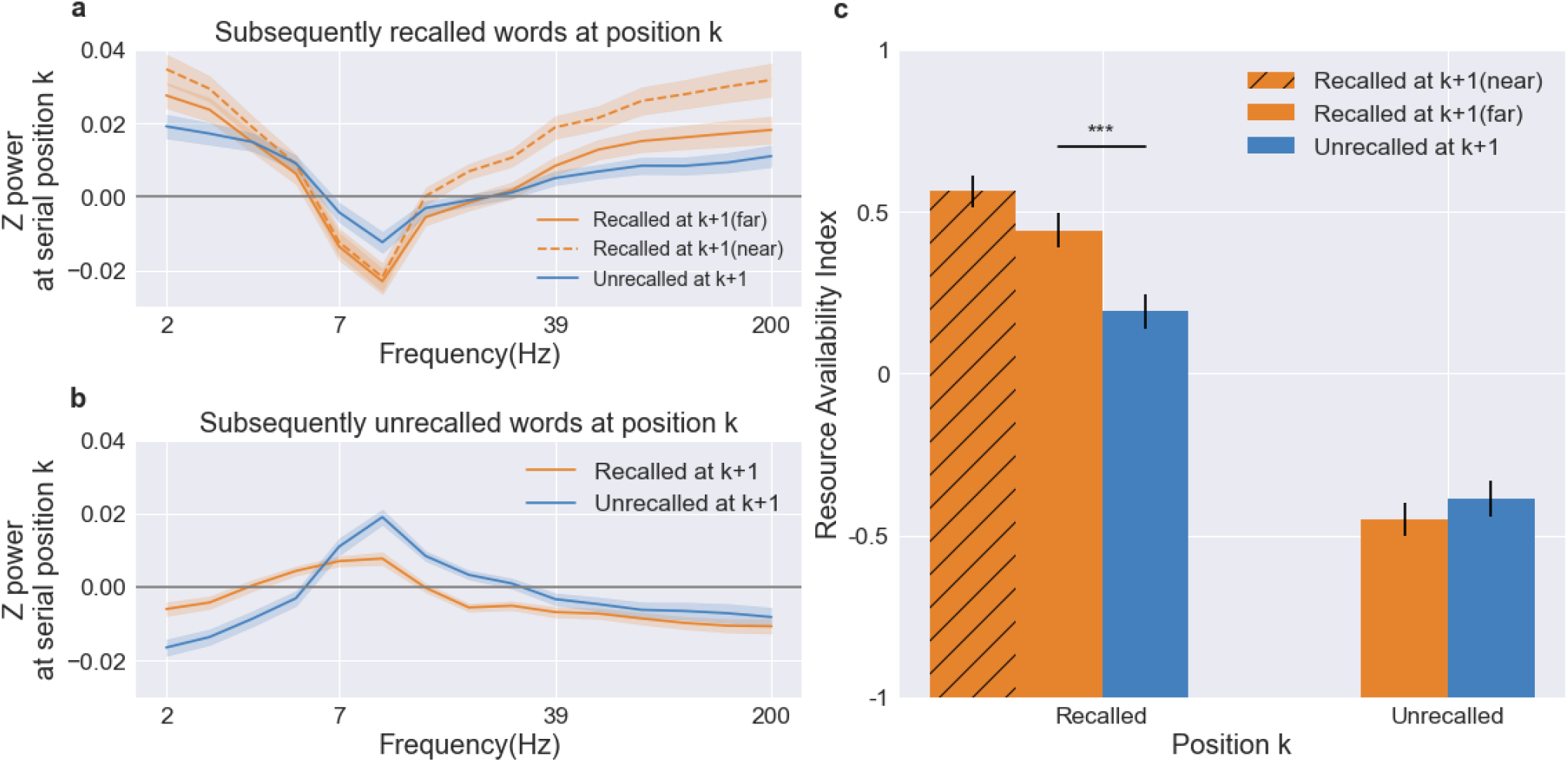
Controlling contiguity effect for sequential subsequent memory effect. a. Different power spectrum patterns when encoding recalled words at position k for subsequently recalled versus unrecalled items encoded at position k+1, further separated into “far” and “near” conditions. b. Different power spectrum patterns when encoding unrecalled words at position k for subsequently recalled versus unrecalled items encoded at position k+1, conditioned on when items at position k were subsequently unrecalled. c. The neural index of resource availability when encoding items at position k, conditioned on whether items at position k and k+1 were recalled together, recalled separately or unrecalled. Asterisks indicate a significant difference between groups (***p *<* .001).

## Appendix D Per-participant level model fitting and results

The SAC model has always been fitted to averaged group data via Monte Carlo simulations (Popov & Reder, 2020), and we followed this tradition when obtaining the group-level model. Specifically, calculating the resources available at each trial requires knowing the resources available on each trial before that, which depends on the specific sequence of word frequencies. At the group-level fitting, the original model iterates over trials, a process that is time-consuming. However, at the per-participant level, instead of fitting via Monte Carlo simulations we fit by a new analytical solution for the simplified model presented in this paper (detailed in Appendix E). This much faster model fitting process given by the analytical solution makes it possible for us to fit the model to each individual subject. Given the limited data available for each participants, we reduce the complexity of the model by only fitting four out of five of the model parameters. We assume that only the depletion rate *δ*, recovery rate *r*, retrieval threshold *θ*_*epi*_, and retrieval noise *σ*_*epi*_ vary among subjects, while the scaling parameter *τ* remains constant across subject which is set as the estimated group-level parameter (*τ* = 0.2). The model was fit to each participant using maximum likelihood 400 times with different starting parameters. 32 participants were excluded because we could not uniquely identify a single set of parameters for them.

Fig. A4 and A5 plot the fits of models at the participant level. First, despite the large variability in the serial position curves of individual subjects (e.g. steepness of the primacy gradient, the location of the asymptote before the recency effect appears, overall levels of recall), the model provided striking fits to each subject’s primacy function (see Fig. A4). Therefore, modeling the primacy effect as resulting from resource depletion and recovery is able to explain not only the aggregated primacy effect, but also individual differences in it (with the exception of one subject who showed no primacy effect).

**Fig.A4.**
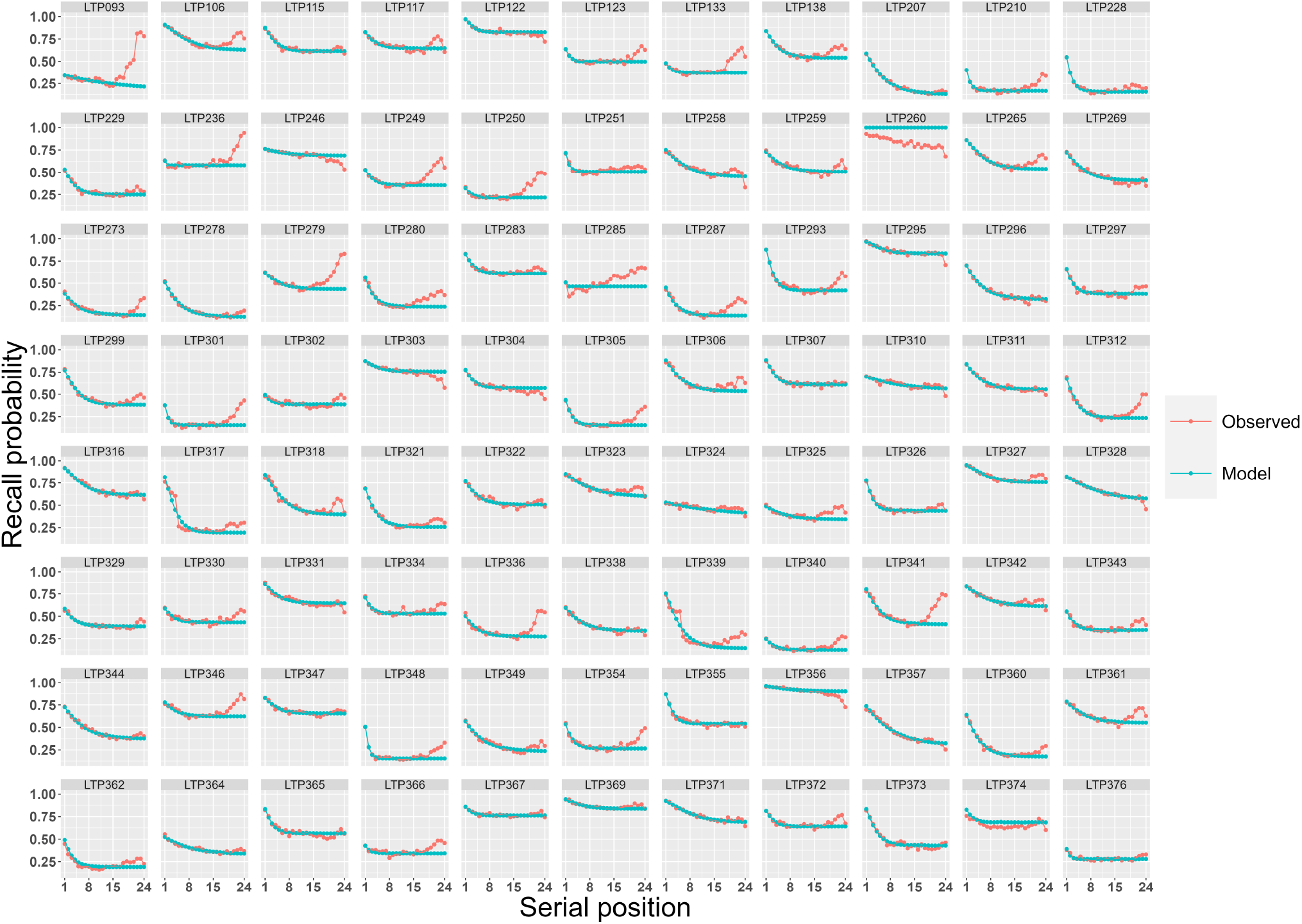
Primacy serial position effect for each participant captured by the SAC model.

Furthermore, the model also provides a good fit to individual differences in the magnitude of the sequential frequency effect (see Fig. A5, mean recall level removed to highlight slope). To estimate numerically how well the model fits the slope of the sequential frequency effect at the level of individual participants, we fit two mixed regression models: 1) regressing the observed recall probability as a function of prior item frequency, with random slopes for each participant, and 2) the same but for the model predicted probability as a dependent variable. Then, we calculated the correlation between the random subject slopes in the two regression model: *r*(86) = 0.57, *p <* 0.001. This result indicates that the model does a decent job at capturing individual differences in the strength of the sequential frequency effect, just by varying the resource depletion and recovery parameters across participants. The distribution of individual parameter values is shown in Fig. A6.

**Fig.A5.**
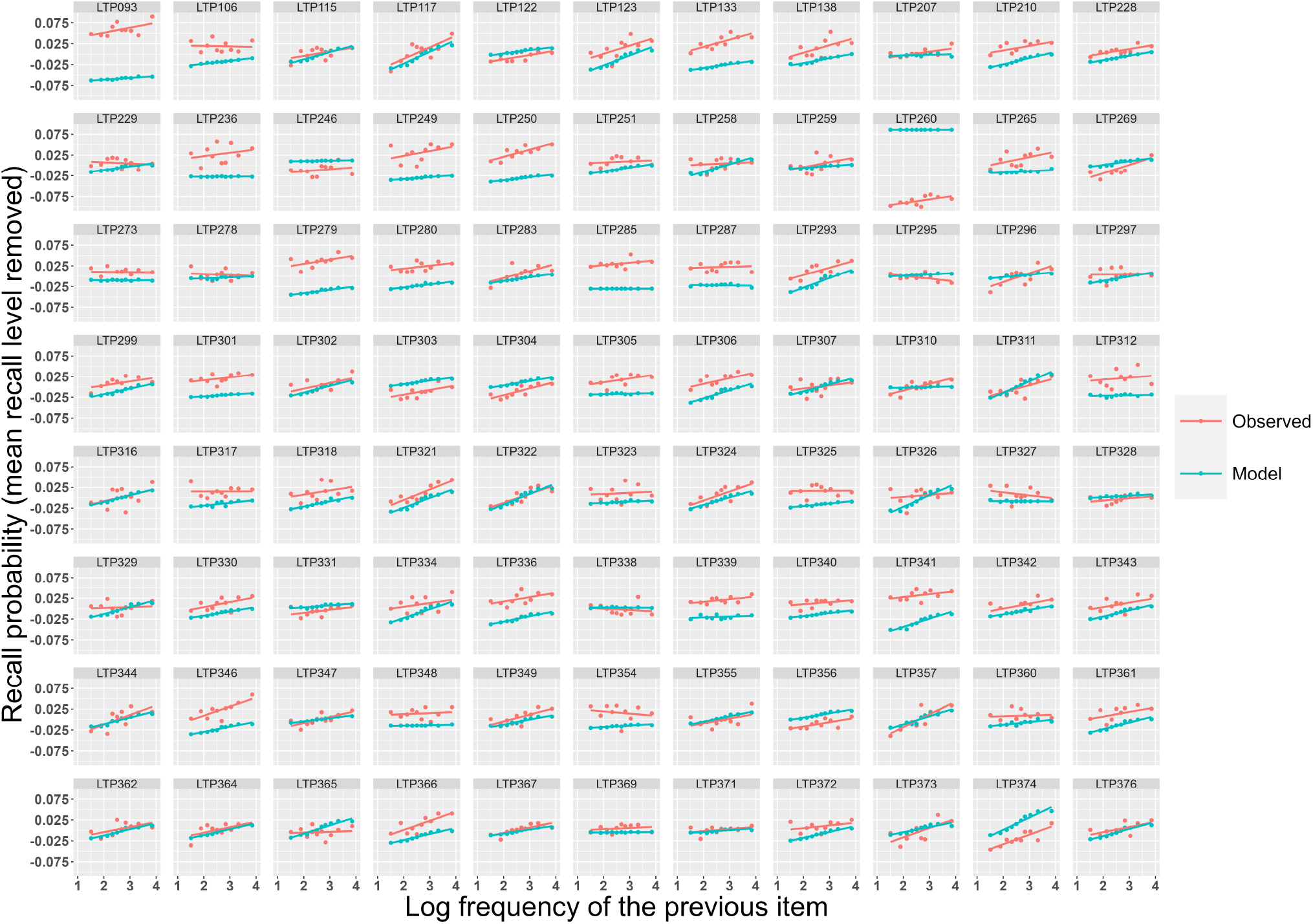
Sequential frequency effect for each participant captured by the SAC model.

**Fig.A6.**
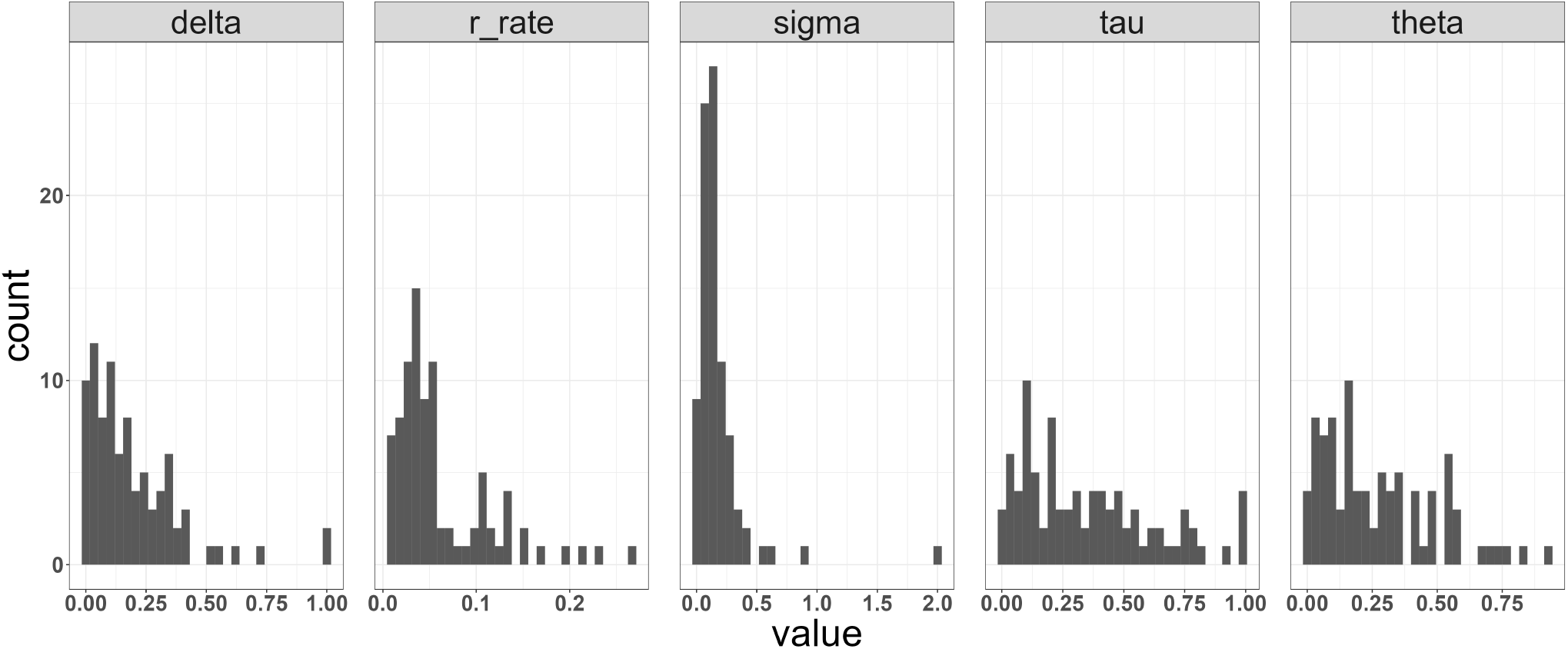
Distribution of individual SAC model parameters - delta (*δ*), r rate (*r*), sigma (*σ*_*epi*_),tau (*τ*) and theta (*θ*_*epi*_).

## Appendix E Derivation of model equations

Here we show how to derive Equation 6 in the main text from equations 1-5. Let *W*_*i*_ denote the amount of available resources at the beginning of the i-th position in a study list. According to Equation 5 from the main text (ignoring the fact that resources cannot exceed a fixed maximum):

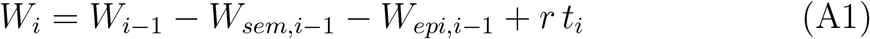

Where

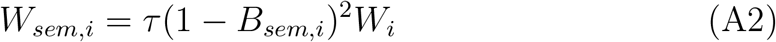

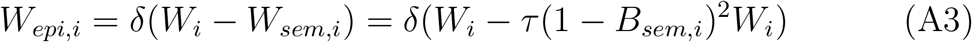

substituting the terms from Eq. A1 with the ones from Eqs. A2-3, we get:

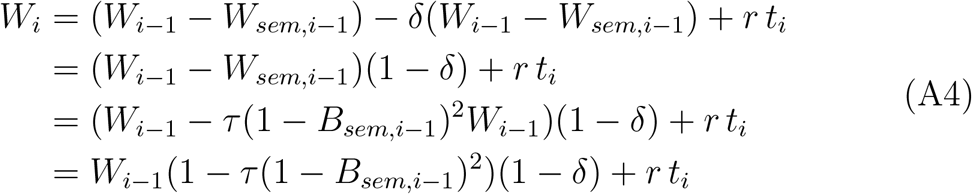

Equation A4 specifies how to calculate *W*_*i*_ based on the resources available on the previous trial and the model parameters, but still requires us to loop over trials in order to calculate the resources for each. By unpacking Equation A4, we can arrive at an equation for *W*_*i*_ that only depends on the initial amount of resources and model parameters. For convenieces, let us use the following substituion symbols (assuming for simplicity constant ISI):

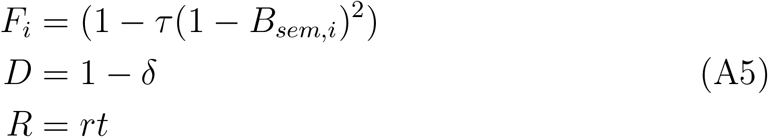

Then Equation A4 becomes:

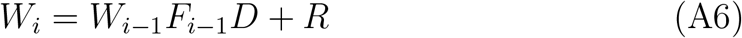

Using Equation A6, we can write out the resources for several sequential positions starting form 1 onwards like this:

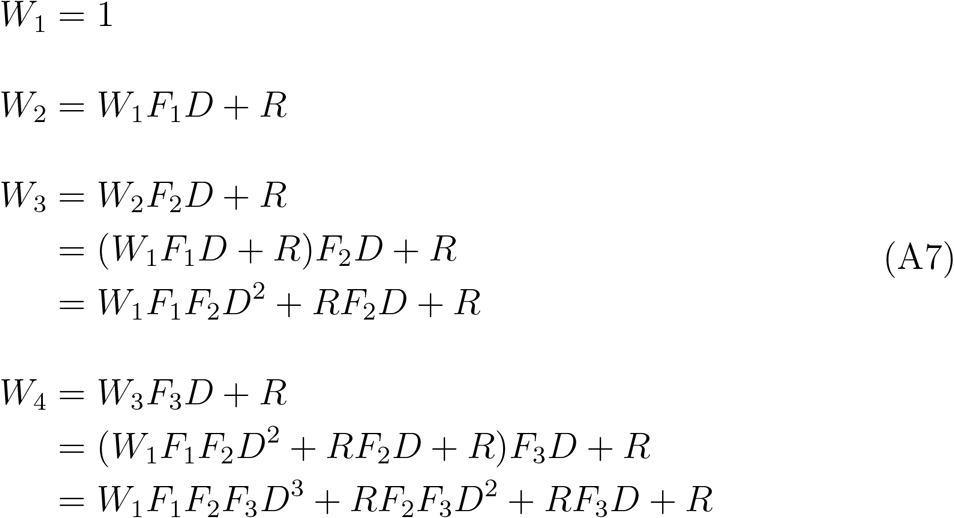

The pattern above is clear. It can be shown that this generalizes to:

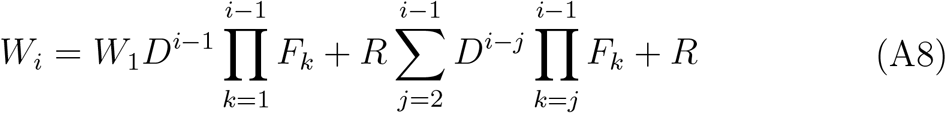

When the initial resources are fixed to 1, as in our simulation, and we expand the substituted terms we get:

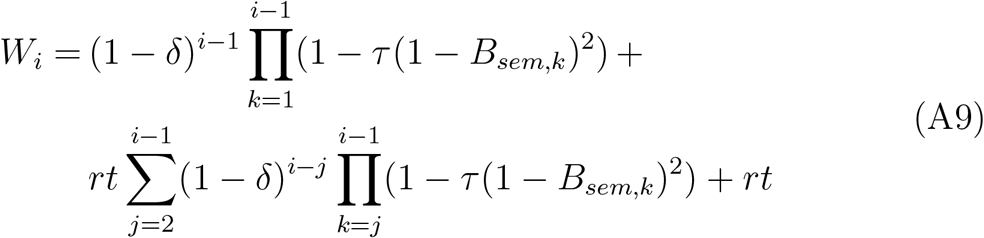

In Equation A9, the available resources on trial i, *W*_*i*_, no longer requires calculating first the resources available on preceding trials. It depends only on the model parameters and the sequence of prior base-level strength of the semantic nodes *B*_*sem*_ for all studied words until trial i.

